# Systematic characterization of therapeutic vulnerabilities in Multiple Myeloma with Amp1q reveals increased sensitivity to the combination of MCL1 and PI3K inhibitors

**DOI:** 10.1101/2023.08.01.551480

**Authors:** Romanos Sklavenitis-Pistofidis, Elizabeth D. Lightbody, Mairead Reidy, Junko Tsuji, Michelle P. Aranha, Daniel Heilpern-Mallory, Daisy Huynh, Stephen J. F. Chong, Liam Hackett, Nicholas J. Haradhvala, Ting Wu, Nang K. Su, Brianna Berrios, Jean-Baptiste Alberge, Ankit Dutta, Matthew S. Davids, Maria Papaioannou, Gad Getz, Irene M. Ghobrial, Salomon Manier

## Abstract

The development of targeted therapy for patients with Multiple Myeloma (MM) is hampered by the low frequency of actionable genetic abnormalities. Gain or amplification of chr1q (Amp1q) is the most frequent arm-level copy number gain in patients with MM, and it is associated with higher risk of progression and death despite recent advances in therapeutics. Thus, developing targeted therapy for patients with MM and Amp1q stands to benefit a large portion of patients in need of more effective management. Here, we employed large-scale dependency screens and drug screens to systematically characterize the therapeutic vulnerabilities of MM with Amp1q and showed increased sensitivity to the combination of MCL1 and PI3K inhibitors. Using single-cell RNA sequencing, we compared subclones with and without Amp1q within the same patient tumors and showed that Amp1q is associated with higher levels of *MCL1* and the PI3K pathway. Furthermore, by isolating isogenic clones with different copy number for part of the chr1q arm, we showed increased sensitivity to MCL1 and PI3K inhibitors with arm-level gain. Lastly, we demonstrated synergy between MCL1 and PI3K inhibitors and dissected their mechanism of action in MM with Amp1q.

## INTRODUCTION

Multiple myeloma (MM) is a bone marrow plasma cell malignancy, whose pronounced genetic heterogeneity prohibits the development of broadly effective precision therapy^1–12^. The most common mutations, affecting *KRAS* and *NRAS*, only occur in about 20% of patients, while most driver genes are mutated in few patients^1–12^. On the other hand, approximately half of MM patients carry translocations involving IgH and one of five major partner genes (*CCND1*, *CCND3*, *WHSC1*/*FGFR3*, *cMAF*, *MAFB*), and ∼85% of patients carry copy number abnormalities, some of which are recurrent^1–17^. While therapeutic targeting of the most frequent translocation, t(11;14), with BCL-2 inhibition has shown promising results in clinical trials, no precision therapy strategy exists for patients with recurrent copy number abnormalities^18–21^. This is largely attributable to innate challenges in targeting a copy number event, such as the multitude of genes involved both *in cis* and *in trans*, as well as difficulties in experimentally modeling such events. As the engineering of whole arm deletions has recently become possible, this holds particularly true for arm-level amplification events, which are more difficult to engineer and thus suffer from lack of proper controls^22–25^.

Gain or amplification of the long arm of chromosome 1 (chr1q), which will be collectively referred to as Amp1q hereafter, is the most frequent copy number amplification in patients with multiple myeloma^26–28^. It is present in approximately 40% of patients at diagnosis and approximately 70% of patients at relapse, suggesting a potential role in disease progression and relapse despite recent advances in MM therapeutics^26–40^. Most importantly, though, it is one of the few genetic alterations with an established negative impact on survival, which is proportional to both the number of cells that carry the event, as well as the number of copies, with 4 copies of chr1q being associated with worse outcome compared to 3 copies^1, 26–41^. Furthermore, Amp1q, as measured clinically by Fluorescence In Situ Hybridization (FISH), is used in clinical practice for patient risk stratification, as it confers high risk of progression in patients with Smoldering Multiple Myeloma (SMM) and MM^42–45^. For these reasons, discovering ways to target Amp1q in patients with MM could significantly affect clinical practice and patient outcomes.

Two minimal common regions have been identified on chr1q, 1q21-1q23 and 1q43-1q44. These regions encompass such candidate genes as *MCL1*, *BCL9*, *CKS1B*, *ILF2*, and *PBX1*, which have been shown to confer sensitivity to different inhibitors^46–56^. However, a systematic characterization of the genetic dependencies and therapeutic vulnerabilities of MM with Amp1q has not been performed.

Here, we employed large-scale CRISPR/RNAi screens and drug screens to characterize and target the dependency landscape of 1q-amplified multiple myeloma and identified MCL1 and the PI3K pathway as actionable therapeutic vulnerabilities in MM with Amp1q. To overcome common challenges associated with the lack of isogenic cell lines, we used single-cell RNA sequencing to compare subclones with and without Amp1q within the same patient tumors, and isolated isogenic clones from a MM cell line allowing us to compare focal and arm-level gain in respect to the sensitivity conferred to MCL1 and PI3K inhibition. Lastly, we dissect the mechanism of action of MCL1 and PI3K inhibition and provide insight into their synergistic activity in MM with Amp1q.

## METHODS

### MM cell line selection

Multiple Myeloma cell lines were selected based on their number of chr1q copies, as measured in copies of *CKS1B* (a marker of Amp1q used in Fluorescence In Situ Hybridization (FISH)) by whole exome sequencing as part of the Cancer Cell Line Encyclopedia (CCLE) (https://depmap.org/portal/download/all/, version 2013-12-03, accessed on 4/10/2023). Copy ratio estimates for *CKS1B* (which reflect the estimate of the segment on which *CKS1B* resides) were transformed into absolute copy numbers by raising 2 to the power of the copy ratio and multiplying that by 2. Copy numbers were then rounded to the nearest integer or integer + 0.5 states and cell lines with a copy number of 2 were considered diploid for chr1q. To ensure cell lines with amplifications of segments excluding *CKS1B* were not misclassified as negative for Amp1q, copy number status was confirmed on IGV. All cell lines used for *in vitro* experiments were tested for mycoplasma contamination (ATCC) and subjected to STR fingerprinting (Labcorp).

### Genome-wide CRISPR screen & analysis

MM1S and KMS18 cells were first virally transduced with LentiCas9-Blast (Addgene #52962) and Cas9 expression was confirmed by Western Blot (Cell Signaling Technology #14697). Subsequently, Cas9-positive cells were virally transduced with subpools of the Avana library of single guide RNAs (sgRNAs) in triplicate (∼1K cells/guide), and samples were drawn on day 4 and day 27 following infection. DNA was extracted using QIAGEN DNA micro kits and deep targeted sequencing of the sgRNAs was performed at the Massachusetts General Hospital Center for Computational & Integrative Biology DNA Core. Following deconvolution and barcode quantitation with PoolQ, read counts were normalized for total depth, multiplied by 10^6, and log_2_-transformed with a pseudocount of 1. Replicates with < 200K reads or correlation < 0.7 were discarded. The remaining technical replicates per subpool were collapsed by computing their median. Log_2_ fold-changes were computed using day 4 as the baseline estimate and normalized by subtracting the median log_2_ fold-change of all non-targeting sgRNAs in the respective subpool. A final guide-level matrix was obtained by computing the median log_2_ fold-change across subpools. Copy number-corrected gene-level matrices were obtained using Ceres following z-mad normalization, as described previously, and used for downstream analyses^57^.

### Targeted C911 RNAi screen & analysis

A custom library of 6,500 short hairpin RNAs (shRNAs), comprising 3,250 targeting shRNAs and their corresponding C911 hairpins, was synthesized at the Broad Institute of MIT and Harvard. Four MM cell lines (KMS18, MM1S, NCIH929, OPM2), two breast cancer cell lines (DU4475, MDAMB231), and three lung cancer cell lines (NCIH1437, NCIH1793, NCIH1650) were virally transduced with the hairpin library in six replicates, and samples taken on days 3 and 27 following infection were subjected to deep targeted sequencing at the Genomics Platform of the Broad Institute. Following deconvolution and barcode quantitation, read counts were normalized for total depth, multiplied by 10^6, and log_2_-transformed with a pseudocount of 1. Replicates with correlation < 0.7 and hairpins with median baseline representation < 1 were discarded. Log_2_ fold-changes were computed using day 3 as the baseline estimate and normalized by subtracting the median log_2_ fold-change of all non-targeting hairpins. Replicates were z-mad normalized and collapsed by computing their mean for a final hairpin-level matrix. Gene-level matrices were obtained using DEMETER2 with C911 hairpins provided for adjustment of the seed effect and used for downstream analyses^58^.

### Dependency Map CRISPR/RNAi datasets & dependency analyses

Ceres and DEMETER2-processed CRISPR & RNAi datasets from the Dependency Map (v.18Q4) were accessed on 12/12/2018^59^. The CRISPR dataset contained the MM cell lines below: EJM, JJN3, KMS11, KMS20, KMS26, KMS27, LP1, MM1S. The RNAi dataset contained the MM cell lines below: KMS18, SKMM2, MM1S, OPM2, RPMI8226, OPM1, KMS27.

Genes with a negative dependency score and whose median score in 1q-amplified MM cell lines was more negative compared to non-amplified controls in the Dependency Map datasets and the in-house genome-wide CRISPR screen, were considered for further filtering. CCLE gene expression data (v.18Q2, accessed on 12/12/2018) were used to identify genes whose difference in median expression levels between 1q-amplified and non-amplified MM cell lines (n=22 and n=3, respectively) was > 2 or < -2, which correspond to the ∼95^th^ and 5^th^ quantiles of the distribution, respectively. Differentially essential genes were then filtered for genes with an up- or downregulation in expression & and an upregulation in expression only for those located on chr1q. Genes that hit in at least two datasets (n=1,523) were considered for downstream analyses. Gene set enrichment analysis was performed using hypergeometric tests against the Hallmark pathway collection from MSigDB (https://www.gsea-msigdb.org/gsea/msigdb/human/collections.jsp, accessed on 4/21/2023) and p-values were corrected using the Benjamini-Hochberg approach. Pathways with a q-value < 0.05 (n=14) were considered enriched in MM cell lines with Amp1q.

### Connectivity Map-guided drug screen

A lineage-agnostic Amp1q gene expression signature was derived by performing differential expression analysis between tumors with and without Amp1q across patients with MM (GSE2658), breast cancer, and lung squamous cell carcinoma (TCGA). Breast cancer and lung squamous cell carcinoma were selected as examples of solid malignancies that can present with Amp1q. Differential expression analysis was performed using t-tests and genes were ranked according to their -log_10_(p-value) multiplied by the sign of their log_2_ fold-change. Genes located in chr1q were removed from consideration and gene ranks were averaged across the three cohorts. The top 150 up- and down-regulated genes by average rank were used to query the Connectivity Map (CMAP, clue.io) (Supplemental Table 1)^60^. The top 201 compounds that were predicted to reverse this signature were then tested in a drug screen in 384-well plate-format with 5,000 cells per well and four MM cell lines (Amp1q: KMS11, OPM2, NCIH929; Non-1q: KMS18). All compounds were tested at 10uM final concentration for 48hrs and in duplicate, while cell viability was measured using CellTiter-Glo (Promega). Bortezomib and DMSO-only wells were included as positive and negative controls, respectively, and % viability was computed after subtracting the average viability from negative control wells. Compound target annotation was based on the Drug Repurposing Hub (https://clue.io/repurposing#download-data, version: 3/24/2020) and for compounds without such annotation, known targets were identified in the literature. Hits (n=47) were identified based on a median viability in MM cell lines with Amp1q of less than 75% and a decrease in their median viability of more than 25% compared to that of the Non-1q control. Targets supported by more than 1 compound were tested for enrichment in the hitlist using Fisher’s exact test and p-values were corrected using the Benjamini-Hochberg approach.

### Drug Repurposing screen

A drug screen was performed using the Broad Institute’s Drug Repurposing library of 4,986 compounds that have cleared various stages of clinical testing^61^. Compounds were tested in 384-well plate-format at 10uM final concentration (30nL of 10mM pools, mixed with 30uL of cells) for 48hrs and in duplicate, with 5,000 cells per well and four MM cell lines (Amp1q: KMS12BM, MM1S, NCIH929; Non-1q: KMS18), and cell viability was measured using CellTiter-Glo (Promega). Bortezomib and DMSO-only/No Treatment Control (NTC) wells were included as positive and negative controls, respectively, and % viability was computed after subtracting the average viability from negative control wells. Compound target annotation was based on the Drug Repurposing Hub (https://clue.io/repurposing#download-data, version: 3/24/2020). Hits (n=470) were identified based on a median viability in MM cell lines with Amp1q of less than 75% and a decrease in their median viability of more than 25% compared to that of the Non-1q control. Targets supported by more than 5 compounds were tested for enrichment in the hitlist using Fisher’s exact test and p-values were corrected using the Benjamini-Hochberg approach.

### MCL-1 & PI3K inhibition *in vitro*

Two specific MCL-1 inhibitors, S63845 and AZD5991, along with a PI3K inhibitor, AZD8186, were validated *in vitro* in an expanded panel of cell lines: KMS18, EJM, SKMM2, MM1S, KMS11, KMS12BM, OPM2, NCIH929. Ten thousand cells per well were treated in a 96-well plate-format for 72hrs with a range of concentrations (0-50uM in a series of 20 two-fold dilutions) and cell viability was measured with CellTiter-Glo (Promega). Drug sensitivity was compared by calculating IC_50_ values between MM cell lines using four parameter non-linear regression models and one-way ANOVA analysis for multiple comparisons. Drug combinatorial synergy screening was carried out using one MCL-1 inhibitor (S63845) and one PI3K inhibitor (AZD8186) in two Amp1q cell lines, NCIH929 and MM1S, and two Non-1q cell lines, KMS18 and EJM. Testing was performed in a 96-well plate-format with 10,000 cells per well and using dose-response matrices in a 10×10 grid manner (0-10uM in two-fold dilutions) and cell viability was measured with CellTiter-Glo (Promega). The dose-response matrices were then used to compute synergy and antagonism scores using Bliss synergy modelling with Combenefit software.

### Single-cell RNA and B cell receptor (BCR)-sequencing of patient samples with subclonal Amp1q

Two patients with Monoclonal Gammopathy of Undetermined Significance (MGUS) and three patients with Smoldering Multiple Myeloma (SMM) who had subclonal Amp1q by Fluorescence In Situ Hybridization (FISH) were selected for this cohort from the PCROWD study (IRB #14-174). All patients provided informed consent. The review boards of all participating centers approved the study in accordance with the Declaration of Helsinki and the International Conference of Harmonization Guideline for Good Clinical Practice. In total, 6 samples, including 4 bone marrow (BM) and 2 peripheral blood (PB) samples, were collected. Mononuclear cells were ficoll-purified, CD138+ plasma cells were selected using magnetic beads (Miltenyi Biotec) and viably frozen in FBS/10% DMSO. Cell-barcoded libraries were prepared using Chromium Single Cell 5’ and V(D)J Enrichment kits by 10X Genomics and sequenced on NovaSeq 6000 S4 flow cells at the Genomics Platform of the Broad Institute of MIT and Harvard (Cambridge, MA). FASTQ files were demultiplexed and processed using Cell Ranger v.6.0.1 and downstream analyses were conducted using Scanpy v.1.8.2^62, 63^. Droplets with > 15% mitochondrial transcripts, < 200 or > 5,000 genes/cell, and < 400 or > 50,000 UMIs/cell were discarded. Malignant plasma cells were differentiated from normal on a per-sample basis using the matched BCR-seq data, and malignant cells with Amp1q were identified by Numbat^64^. Gene expression levels were compared between clones with and without Amp1q within each tumor using Wilcoxon rank-sum tests and p-values were adjusted with the Benjamini-Hochberg approach. The enrichment of the PI3K pathway in clones with Amp1q was tested by Gene Set Enrichment Analysis (GSEA) within each tumor using the PI3K/Akt/mTOR Hallmark Pathway.

### Single-cell RNA-sequencing of MM cell lines under treatment with MCL1i, PI3Ki or combination

Two MM cell lines with Amp1q (KMS12BM, KMS11) were selected for this experiment. The cells were treated for 6hrs with MCL1i (S63845) at 0.1uM, PI3Ki (AZD8186) at 5uM, their combination, and 0.25% DMSO, and following a wash with PBS/0.04% BSA, viable cells were loaded on the Chromium Controller for encapsulation. Cell-barcoded libraries were prepared using Chromium Single Cell 5’ and V(D)J Enrichment kits by 10X Genomics and sequenced on NovaSeq 6000 S4 flow cells at the Genomics Platform of the Broad Institute of MIT and Harvard. FASTQ files were demultiplexed and processed using Cell Ranger v.6.0.1 and downstream analyses were conducted using Seurat v.4.0.1^62, 65^. Droplets with > 15% mitochondrial transcripts, <2,000 or >6,500 genes/cell, <5,000 or >30,000 UMIs/cell, and < 18,000 reads/cell were discarded, and 18,000 reads per cell were downsampled across the entire cohort. Following downsampling, droplets with > 10% mitochondrial transcripts, and < 3,500 or > 5,000 genes/cell were further discarded. Cell cycle scores were computed using Seurat’s CellCycleScoring() function and custom thresholds were determined empirically for each cell line to assign cells into phases. Specifically, cells were assigned to the S phase if their S score was > 0 for either cell line and less than or equal to 0.5 times the cell’s G2M score, and the G2M phase if their G2M score was > -0.2 for KMS12BM and > -0.5 for KMS11. For each sample, the number of cells in each phase was divided by the total number of cells to compute a proportion and a 95% confidence interval was computed by iteratively subsampling 70% of cells 10,000 times. Empirical p-values were computed for the difference in cell cycle phase proportions between conditions by bootstrapping with 10,000 iterations. Differential expression was performed using Wilcoxon’s rank-sum tests and p-values were adjusted with the Benjamini-Hochberg approach.

### Isogenic clones of KMS12BM with different copy number for 1q31-1q44

Single cells from the MM cell line KMS12BM were sorted (BD LSR Fortessa) into a 96-well plate with 100uL of RPMI 1640/50% FBS per well and cultured for approximately 1 month. Eight daughter clones were established in total. DNA was extracted using QIAGEN DNA micro kit and fragmented targeting 350bp fragments using Covaris M220 and microTUBE-50 tubes (55uL volume, 20C, Peak incident power: 75W, Duty factor: 10%, Cycles per burst: 200, Duration: 80s). Genomic libraries were prepared using KAPA HyperPrep kits (Roche), following the protocol for PCR-free whole-genome sequencing (WGS), and sequenced at ∼8.5X depth on a NovaSeq 6000 SP flow cell with 300 cycles (150bp reads) at the Genomics Platform of the Broad Institute of MIT and Harvard. FASTQ files were demultiplexed using bcl2fastq and aligned to hg19 using bwa-mem. The Genome Analysis Tool Kit (GATK) somatic copy number variant (CNV) analysis workflow (v.4.2.0.0) with a changepoint penalty factor of 50 and a custom CNV panel of normal (PoN) were used to characterize the copy number profile of the isolated clones^66^. Two clones, one with 4 copies of chr1q (Clone 1), and one with 4 copies of 1q21-1q25 and 3 copies of 1q31-1q44 (Clone 2), were treated *in vitro* with MCL1i (S63845) and PI3Ki (AZD8186) using a range of 20 concentrations (0.0001-20uM). Cell viability was measured using CellTiter-Glo (Promega) and compared with one-way Anova and Tukey’s test.

### BH3 profiling of MM cell lines & KMS11 following PI3K inhibition

Three million cells per sample/condition were harvested, centrifuged and resuspended with 550uL of MEB2P buffer (150 mM mannitol, 10 mM HEPES-KOH pH 7.5, 150 mM KCI, 1 mM EGTA, 1 mM EDTA, 0.1% BSA, 5 mM succinate, 0.25% poloxamer 188) prior to adding 15uL to each well of a 384-well plate pre-made with individual BH3 peptides (BIM, BAD, PUMA, HRKy, MS1, New England Peptides, Inc) or BH3 mimetics (ABT199, #S8048, Selleckchem, A1331852, #S7801, Selleckchem) at various concentrations and 0.002% digitonin, dissolved in 15uL of MEB2 buffer. Cells were incubated for 1 hour at room temperature (RT) to allow cell membrane permeabilization by digitonin and BH3 peptide/mimetic entry to induce cytochrome *c* loss. Subsequently, 15uL of 4% paraformaldehyde (PFA) were added to each well for 30 minutes at RT for fixation, followed by 15uL of N2 buffer (1.7 M Tris, 1.25 M glycine pH 9.1) for 20 minutes at RT to neutralize the PFA. Next, 10uL of staining cocktail containing 1x Tween20, anti–CytC–Alexa Fluor 488 (#612308, Biolegend) and Hoechst-33342 (#H3570, Invitrogen) were added to each well and incubated overnight prior to reading on BD FACS Fortessa and analyzing using the FlowJo software. Dynamic BH3 profiling was performed after treating cells with the PI3K inhibitor AZD8186 at a final concentration of 5uM.

## RESULTS

### Systematic characterization of genetic dependencies in Multiple Myeloma with Amp1q

We set out to systematically map genetic dependencies associated with Amp1q in MM. Due to the technical challenges associated with generating isogenic cell lines for a chromosomal amplification event and the rarity of MM cell lines without Amp1q, the discovery of genetic dependencies requires the comparison of non-isogenic cell lines with and without Amp1q^25, 28^. Such comparisons are inherently noisy due to inter-tumor variability that is unrelated to Amp1q, as well as technical variables associated with the technology used (RNAi vs CRISPR) and the pooled format^67^. To control for technical noise and the low number of controls per dataset, we leveraged both RNAi and CRISPR screen data from the Cancer Dependency Map (DepMap) (Amp1q: n=12; Non-Amp1q: n=3) and performed a genome-wide CRISPR screen on a MM cell line with Amp1q and one without, which allowed us to filter for genes that hit in at least two of the three datasets. To control for noise due to differences unrelated to Amp1q, we filtered for genes whose expression was either up- or down-regulated in MM cell lines with Amp1q, reasoning that Amp1q could result in differential dependencies due to either up- or down-regulation of genes *in trans*. We defined hits as genes with: (i) a negative median dependency score in MM cell lines with Amp1q and a negative difference compared to the median dependency score in MM cell lines without Amp1q, and (ii) a difference in median expression between MM cell lines with and without Amp1q of > 2 or < -2. Overall, 2,101 genes hit in one dataset, 1,243 genes hit in two datasets, and 280 genes hit in all three datasets, without clear technology-based bias (**Figure 1A, B**). Considering genes that hit in at least two datasets, we split each chromosome into 30 equal-sized bins and identified three segments with at least 20 hits and significant positional enrichment (hypergeometric, q < 0.05), with chr1q:149-158Mb (corresponding to cytobands 1q21-1q23) being the segment with the most hits (n=34) (**Figure 1C**). This observation suggests that our filtering strategy enriched for hits associated with Amp1q, with minimal positional bias due to other inter-tumor variability. Next, to identify pathway dependencies related to genome-wide perturbations in MM cell lines with Amp1q, we performed gene set enrichment analysis using hypergeometric tests against the Hallmark pathway collection. We found that pathways related to the cell cycle, DNA repair, metabolic activity, protein translation and degradation, as well as the PI3K-mTOR pathway were essential for cell survival in MM cell lines with Amp1q (**Figure 1D**). For genes residing on chr1q, we further filtered for genes with an upregulation in expression only and observed a positional enrichment for genes located in the 1q21-1q23 region (OR = 1.95, 95% CI: 1.42-2.68, Fisher’s exact, p = 2e-05) (**Figure 1E**). This is consistent with prior studies which identified 1q21-1q23 as a minimal common region in patients with MM and Amp1q^46^. Hits included established targets, such as *MCL1*, *BCL9*, and *ILF2*, as well as novel targets, such as *CLK2*, a kinase involved in RNA splicing, and *PIP5K1A*, a kinase responsible for synthesizing PIP2, the substrate of PI3K kinase. We confirmed the increased dependency of MM cell lines with Amp1q on MCL1 by BH3 profiling, which showed increase cytochrome *c* release with MS1 peptide (an MCL1 inhibitor) treatment in cell lines with Amp1q (**Figure 1F, G**). To validate our novel targets, we performed a targeted shRNA screen targeting genes in the 1q21-1q23 region in four MM cell lines, three with Amp1q and one without (**Figure 1H**). Novel targets such as *CLK2* and *PIP5K1A* were confirmed using this approach. Lastly, to explore the lineage specificity of our hits, we analyzed DepMap CRISPR screen data from lung and breast cancer cell lines with and without Amp1q and performed a targeted shRNA screen on two breast cancer (Amp1q, n=1; Non-Amp1q, n=1) and three lung cancer cell lines (Amp1q, n=2; Non-Amp1q, n=1) (**Figure 1I, J**). Interestingly, *MCL1* and *CLK2* appeared to be lineage-agnostic dependencies in cancer with Amp1q, suggesting that targeted therapy with MCL1 inhibitors (MCL1i) or CLK2 inhibitors (CLK2i) may be appropriate for patients with a variety of 1q-amplified malignancies.

**Figure 1.**
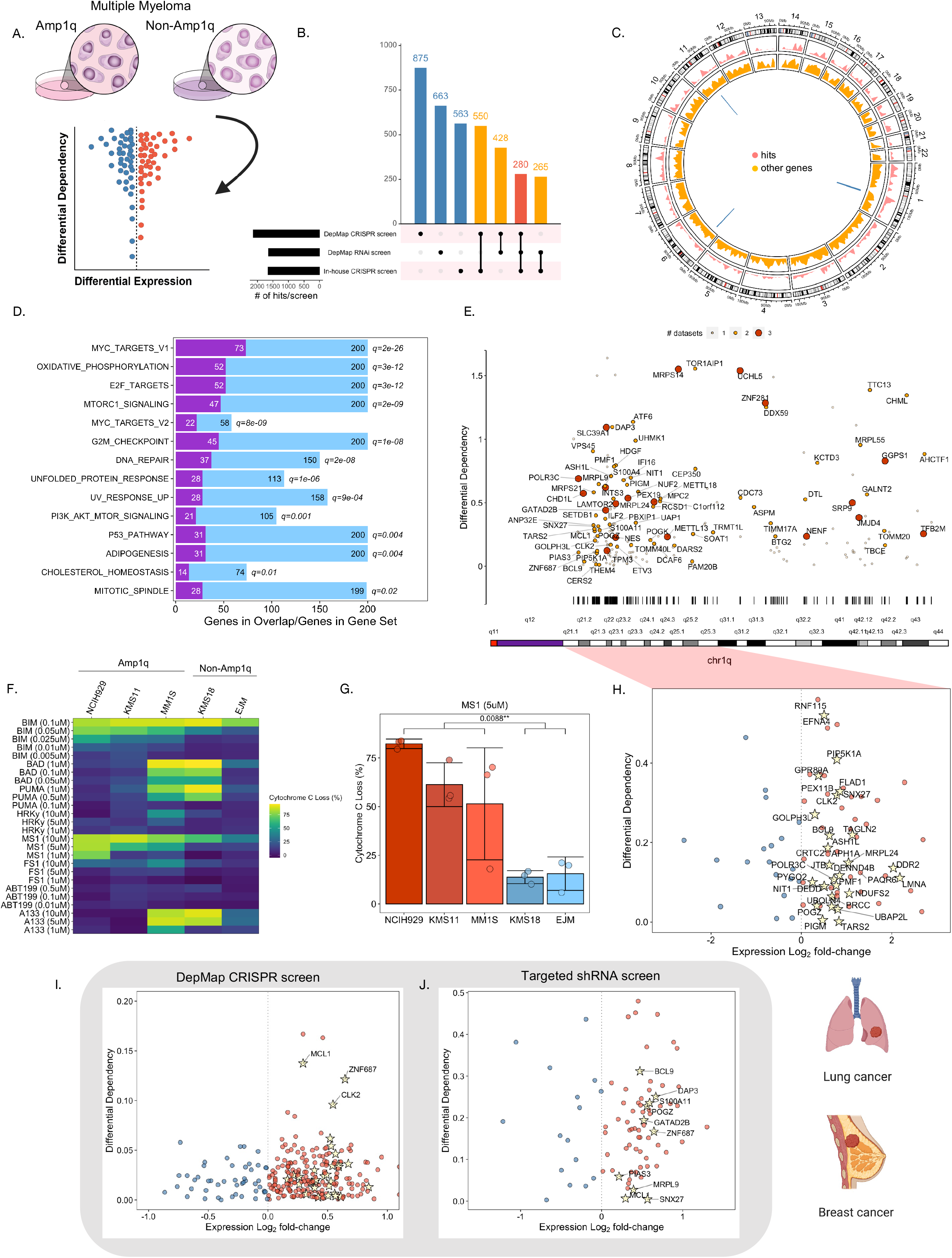
Systematic characterization of genetic dependencies in Multiple Myeloma with Amp1q. A) Schematic of experimental and analytical approach. B) UpSet plot of hits in the dependency analyses by dataset. C) Circos plot of the positional enrichment of genes that hit (red) or did not hit (orange) in the dependency analyses. Blue bars correspond to regions with significant enrichment of hits (hypergeometric, q < 0.05). D) Barplot of gene set enrichment analysis results for pathways with hypergeometric q < 0.05. In light blue, the number of genes in the gene set (black letters); in purple, the number of hits in a given gene set (white letters). E) Scatter plot of hits located on chr1q; genes are colored based on the number of datasets in which they hit (range: 1-3); the positional density of hits along chr1q is visualized with black lines aligned with a chr1q ideogram. F) Heatmap of cytochrome *c* loss (%) in cell lines with (n=3) and without Amp1q (n=2). G) Barplots of cytochrome *c* loss (%) in cell lines with (n=3) and without Amp1q (n=2) incubated with 5uM of the MS-1 peptide. Error bars correspond to the standard deviation across three replicates and the p-value was computed with Wilcoxon’s rank-sum test. H) Scatterplot of log_2_ fold-change in expression between MM cell lines with and without Amp1q (x-axis) and differential dependency (i.e., the difference between median dependency estimates in MM cell lines with and without Amp1q) in the targeted shRNA screen dataset (y-axis). Genes with a positive fold-change are colored in red, while genes with a negative fold-change are colored in blue. Genes that hit in both the targeted shRNA dataset and the genome-wide screens are marked with yellow stars. I) Scatterplot of log_2_ fold-change in expression between breast/lung cancer cell lines with and without Amp1q (x-axis) and differential dependency (i.e., the difference between median dependency estimates in breast/lung cancer cell lines with and without Amp1q) in the DepMap CRISPR screen dataset (y-axis). Genes with a positive fold-change are colored in red, while genes with a negative fold-change are colored in blue. Genes that hit in both this dataset and the MM genome-wide screens are marked with yellow stars. J) Scatterplot of log_2_ fold-change in expression between breast/lung cancer cell lines with and without Amp1q (x-axis) and differential dependency (i.e., the difference between median dependency estimates in breast/lung cancer cell lines with and without Amp1q) in the targeted shRNA screen dataset (y-axis). Genes with a positive fold-change are colored in red, while genes with a negative fold-change are colored in blue. Genes that hit in both this dataset and the MM genome-wide screens are marked with yellow stars.

### Two large-scale drug screens identify the PI3K-mTOR pathway as a therapeutic vulnerability in Multiple Myeloma with Amp1q

Genetic dependencies do not always translate into therapeutic options, due to issues related to target druggability, the specificity and effectiveness of available compounds, and functional redundancy. To systematically characterize the drug sensitivities of MM with Amp1q, we performed two drug screens: (i) a drug repurposing screen of 4,986 compounds that have cleared various stages of clinical testing (Amp1q, n=3; Non-Amp1q, n=1), and (ii) a screen of 201 compounds predicted to reverse a lineage-agnostic gene expression signature associated with Amp1q by the Connectivity Map (CMap) (Amp1q, n=3; Non-Amp1q, n=1) (Methods). Both screens were conducted in duplicate using compounds at 10uM concentration for 48hrs. In both screens, hits were identified based on a median viability in MM cell lines with Amp1q of less than 75% and a decrease in their median viability of more than 25% compared to that of the Non-Amp1q control. As expected, most compounds in the drug repurposing screen were inactive, while 470 compounds were selected as hits (**Figure 2A**). Drug targets with > 5 compounds in our hit list were tested for enrichment using Fisher’s exact tests, which revealed PI3K-mTOR (q=4e-16) and HDAC inhibitors (q=4e-14) as the top drug sensitivities in MM with Amp1q (**Figure 2B, C**). In contrast to the drug repurposing screen, compounds used in the CMap-guided screen were mostly effective, and 47 compounds were selected as hits (**Figure 2D**). Drug targets with > 1 compound in our hit list were tested for enrichment using Fisher’s exact tests. Once again, PI3K-mTOR inhibitors were significantly enriched for hits (q=0.001) (**Figure 2E, F**). These results are consistent with our dependency analyses demonstrating increased dependency of MM with Amp1q on the PI3K-mTOR pathway and suggest that PI3K inhibitors (PI3Ki) may represent a therapeutic option for MM with Amp1q.

**Figure 2.**
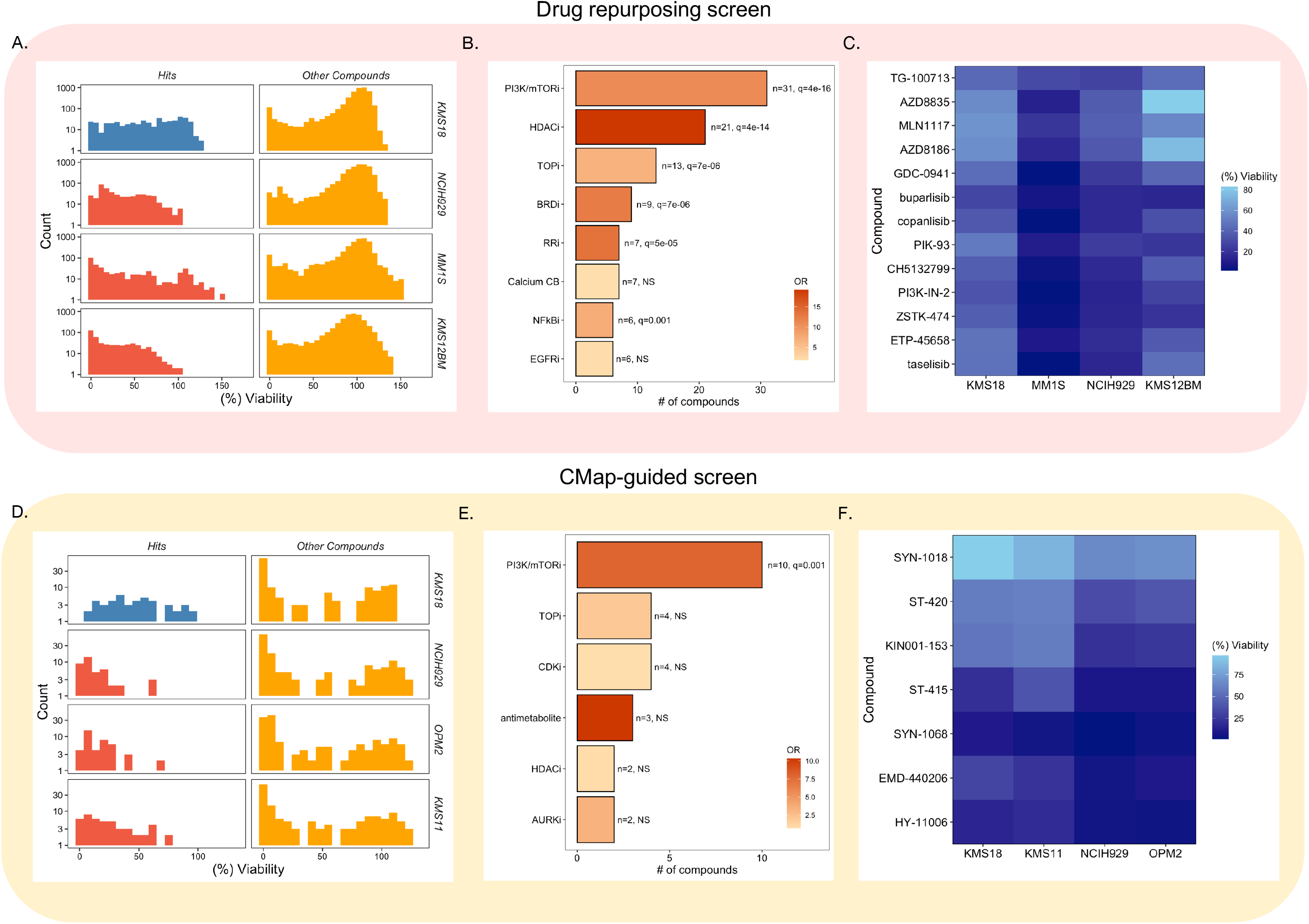
Two large-scale drug screens identify the PI3K-mTOR pathway as a therapeutic vulnerability in Multiple Myeloma with Amp1q. A) Histogram of (%) cell viability with compounds that hit (blue: non-Amp1q; red: Amp1q) or other compounds in the drug repurposing screen across MM cell lines. B) Barplot of compound target enrichment analysis results from the drug repurposing screen. Targets are sorted in decreasing order of compound hits. Targets with >5 compounds in the hit list were tested for enrichment using Fisher’s exact test. C) Heatmap of (%) cell viability across MM cell lines with PI3K inhibitors that hit in the drug repurposing screen. D) Histogram of (%) cell viability with compounds that hit (blue: non-Amp1q; red: Amp1q) or other compounds in the CMap-guided drug screen across MM cell lines. E) Barplot of compound target enrichment analysis results from the CMap-guided drug screen. Targets are sorted in decreasing order of compound hits. Targets with >1 compound in the hit list were tested for enrichment using Fisher’s exact test. F) Heatmap of (%) cell viability across MM cell lines with PI3K inhibitors that hit in the CMap-guided drug screen.

### In vitro validation and synergy of MCL1 and PI3K inhibitors in MM cell lines with Amp1q

To confirm *MCL1* and the PI3K pathway as therapeutic vulnerabilities in MM with Amp1q, we tested two *MCL1* inhibitors and a PI3K inhibitor in an expanded panel of 8 MM cell lines (Amp1q: n=5; Non-Amp1q: n=3). We found that both *MCL1* inhibitors and the PI3K inhibitor were more effective in MM cell lines with Amp1q compared to those without (**Figure 3A**). All five Amp1q cell lines showed moderate to high sensitivity to AZD5991 (0.016-0.39μM) and high sensitivity to S63845 (0.0034-0.1μM), while non-Amp1q lines showed moderate sensitivity to insensitivity to both AZD5991 (0.56-28.80μM) and S63845 (0.34-27.03μM). The PI3K inhibitor also showed higher activity in Amp1q cells (1.14-8.89μM) compared to non-Amp1q cells (13.47-34.72μM). We observed increased MCL1 protein expression levels with increasing concentration of the MCL1i S63845 (due to the inhibitor outcompeting MCL1 degraders for binding) and decreased pAkt levels with increasing concentration of the PI3Ki AZD8186 in both Amp1q and non-Amp1q cell lines (**Figure 3B**). In other malignancies, *MCL1* may interact with the PI3K pathway^68–72^, therefore, we hypothesized that the combination of MCL1i and PI3Ki may be synergistic in MM with Amp1q. Indeed, we observed synergy in MM cell lines with Amp1q, where the combination of the two drugs led to reduced concentrations required to induce MM cell death for both compounds (**Figure 3C**). At the same concentration in non-Amp1q cells no synergy or additive effects were observed, and synergy was only observed at higher concentrations in the KMS18 non-Amp1q cell line (**Figure 3C**). At the lowest combinatorial treatment doses tested in both Amp1q and non-Amp1q cell lines (0.02μM MCL1i and 1.28μM PI3Ki), combination treatment achieved on average 32-66% more killing in MM cell lines with Amp1q compared to non-Amp1q cell lines (p=0.01) (**Figure 3D**). This is important, as clinical trials of MCL1 inhibitors were halted due to significant toxicity at the target therapeutic doses, and the synergistic combination of MCL1i and PI3Ki may reduce the doses required to achieve responses, resulting in less toxicity and enabling the use of an otherwise potent drug. Furthermore, both types of inhibitors have been clinically tested before, which could accelerate the translation of these findings.

**Figure 3.**
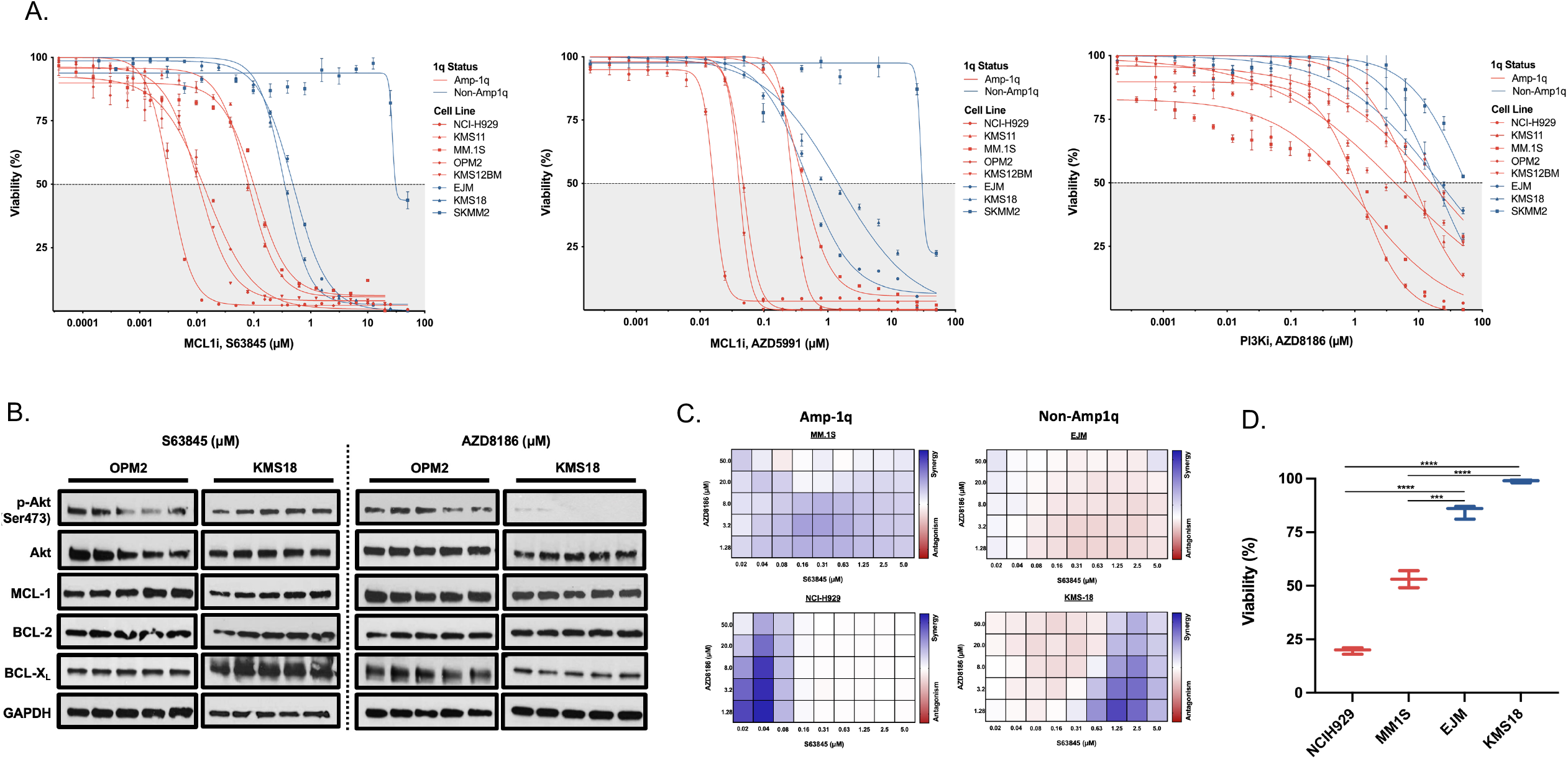
In vitro validation and synergy of MCL1 and PI3K inhibitors in MM cell lines with Amp1q. A) IC_50_ curves of panel of MM cell lines treated with AZD5991, S63845 (MCL1 inhibitors), and AZD8186 (PI3K inhibitor), where red represent cell lines with Amp1q and blue represent non-Amp1q cell lines. B) Western Blot of Amp1q and non-Amp1q cells treated with increasing doses of S63845 and AZD8186 and probed for p-Akt, Akt and various anti-apoptotic protein family members (BCL2, BCL-xL, MCL1), as well as loading control (GAPDH). C) Synergy plots for NCIH929 and MM1S (Amp1q) and EJM and KMS18 (non-Amp1q) treated with 10×10 dose matrices of S63845 and AZD8186, where blue represents synergy and red represents antagonism. D) Viability of NCIH929 and MM1S (Amp1q cells) and EJM and KMS18 (non-Amp1q cells) treated with a combination of 0.02μM MCL1i and 1.28μM PI3Ki, representing the lowest dual concentrations achieving synergy in Amp1q cells.

### Single-cell RNA sequencing of patients with subclonal Amp1q demonstrate association of MCL1 and PI3K dependency with Amp1q

An important limitation imposed by the lack of isogenic cell lines is that the dependencies and therapeutic vulnerabilities discovered may contain targets not associated with the presence of Amp1q. Filtering for targets that are differentially expressed in MM with Amp1q can only partly address this, due to noise imparted by inter-tumor and interindividual variability in differential expression analyses, too^67^. Therefore, we reasoned that differential expression analysis comparing subclones with and without Amp1q within the same tumors could confirm the association of nominated targets with the presence of Amp1q, as it inherently controls for unwanted sources of variability. Subclonal Amp1q is more frequently observed in early-stage disease, such as Monoclonal Gammopathy of Undetermined Significance (MGUS) and Smoldering MM (SMM)^28^. Thus, we performed single-cell RNA sequencing on 6 bone marrow (n=4) and circulating tumor cell (n=2) samples from 5 patients with MGUS (n=2) and SMM (n=3) who had subclonal Amp1q by FISH. In total, 19,818 plasma cells were detected, including 15,194 malignant and 4,624 normal plasma cells (**Figure 4A**). All tumors had subclonal Amp1q, as confirmed by copy number variant inference (**Figure 4B**). In all but one patient, the subclone carrying Amp1q had no other differences in copy number compared to the parental clone; in one patient (P4), the subclone carrying Amp1q also had gains of chr2p and chr4q, as well as Del8p, while the subclone without Amp1q had Del17p (**Figure 4C**). By comparing *MCL1* levels between subclones with and without Amp1q, we found that *MCL1* is consistently overexpressed in the presence of Amp1q (**Figure 4D**). Furthermore, we observed consistent upregulation of the PI3K-mTOR pathway in subclones with Amp1q (**Figure 4E, F**). These results suggest that differences observed in dependency on *MCL1* and the PI3K-mTOR pathway in MM cell lines with Amp1q may indeed be associated with the presence of Amp1q. Furthermore, by systematically looking for differentially expressed genes between subclones with and without Amp1q, we were able to identify frequently dysregulated genes (in at least 3 of the 6 samples) that do not reside on chr1q. We then used this list to filter our non-chr1q dependency hits and derived a short list of 99 genes that are differentially essential and differentially expressed in MM with Amp1q due to trans-regulation (**Figure 4G**). This short list includes such targets as *CCT7*, a chaperone important for protein folding, and *HSPA9*, which encodes a member of the Hsp70 heat shock protein family. Interestingly, the PI3K-mTOR pathway was the top hit in an enrichment analysis, suggesting that the observed dependency on PI3K in MM with Amp1q is not necessarily cis-regulated (**Figure 4H**).

**Figure 4.**
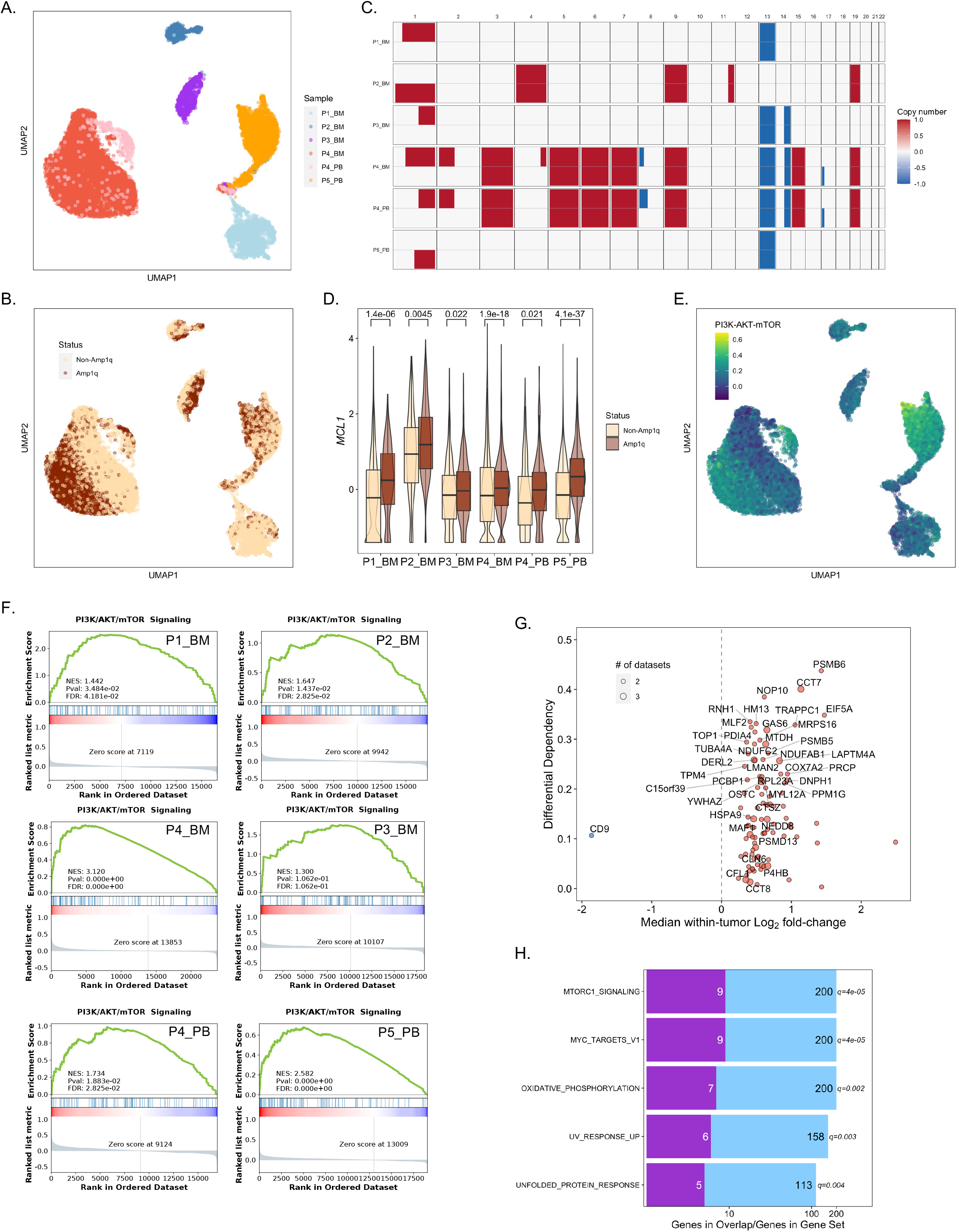
Single-cell RNA sequencing of patients with subclonal Amp1q demonstrate association of MCL1 and PI3K dependency with Amp1q. A) UMAP embedding of malignant plasma cells in the patient cohort. B) UMAP embedding of malignant plasma cells colored based on the copy number status for chr1q (red: Amp1q; beige: non-Amp1q). C) Heatmap of copy number changes in malignant plasma cells of the patient cohort. Each sample is split into two subclones, one with and one without Amp1q. D) Boxplots of *MCL1* expression in subclones with and without Amp1q across all samples. P-values were computed using Wilcoxon’s rank-sum tests and corrected with the Benjamini-Hochberg approach. E) UMAP embedding of malignant plasma cells colored by the activity of the PI3K/Akt/mTOR Hallmark pathway. F) GSEA results for the PI3K/Akt/mTOR Hallmark pathway in subclones with Amp1q across all six samples. G) Scatterplot of the median log_2_ fold-change in expression between subclones with and without Amp1q (x-axis) and differential dependency (i.e., the difference between median dependency estimates in MM cell lines with and without Amp1q) in the genome-wide MM screens (y-axis). Genes with a positive fold-change are colored in red; genes with a negative fold-change are colored in blue. Genes with a differential dependency of at least 0.2 or genes that hit in all three datasets are labeled. H) Barplot of gene set enrichment analysis results for pathways with hypergeometric q < 0.05. In light blue, the number of genes in the gene set (black letters); in purple, the number of hits in a given gene set (white letters).

### Isolation of isogenic clones with different copy number for 1q31-1q44 highlight role of arm-level amplification in MCL1/PI3K inhibitor sensitivity

Because the generation of isogenic clones for chromosomal amplifications is technically challenging, we instead attempted to isolate isogenic clones from a MM cell line with Amp1q (KMS12BM) by sorting single cells in a 96-well plate, establishing daughter clones, and screening them with whole-genome sequencing (WGS) to search for differences in chr1q copy number (**Figure 5A**). In total, we established 8 clones, 7 of which had 4 copies of chr1q (1q21.1-1q44), while 1 (Clone 2) had 4 copies for 1q21.1-1q25.3, where *MCL1* resides, but 3 copies for 1q31.1-1q44, with minimal other differences (**Figure 5B**). Therefore, we used this model to understand whether the copy number of the 1q31.1-1q44 segment, which contains the second minimal common region in MM with Amp1q (1q43-1q44), impacts response to MCL1i and PI3Ki. This is important as our data suggests that PI3K dependency may be the result of broader effects as opposed to the perturbation of a single gene, and we need to know how to properly measure Amp1q in clinical practice to better select patients for targeted therapy. Interestingly, when we treated Clone 2 and a representative specimen from the rest of the clones, Clone 1, with a range of concentrations of the MCL1i S63845 and the PI3Ki AZD8186, we observed an increase in sensitivity to both types of inhibitors with the increase in copy number for segment 1q31.1-1q44 (i.e., in Clone 1) (**Figure 5C, D**). These results suggest that while the amplification of the 1q21.1-1q25.3 segment alone confers sensitivity to these inhibitors, amplification of the rest of chr1q further increases that sensitivity, and thus selecting patients for treatment based on the copy number of the 1q21-1q25 segment alone, as routinely measured by FISH, may not be appropriate. This would require prospective validation in the context of a clinical trial.

**Figure 5.**
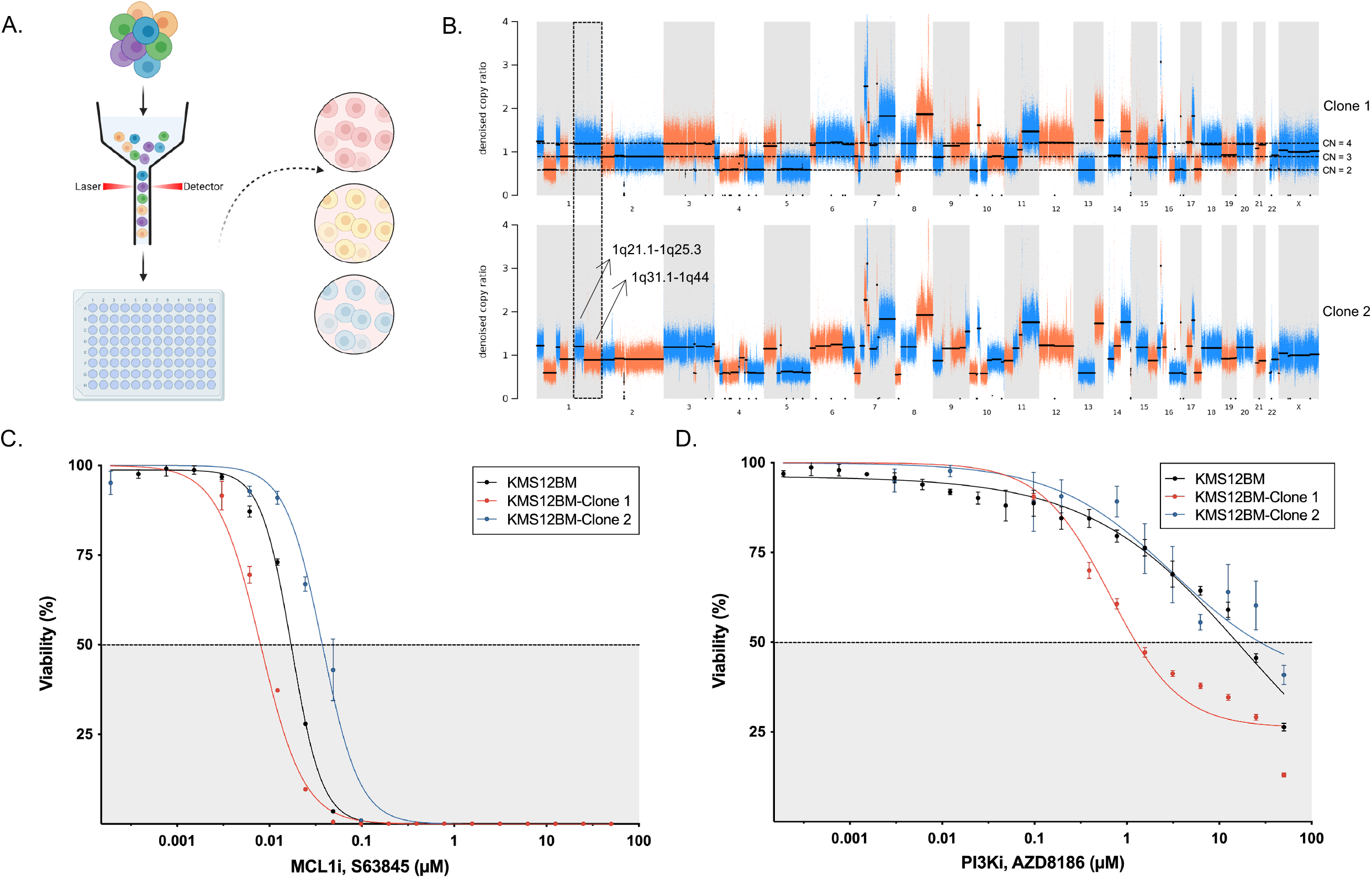
Isolation of isogenic clones with different copy number for 1q31-1q44 highlight role of arm-level amplification in MCL1/PI3K inhibitor sensitivity. A) Schematic of the experimental approach for the isolation of isogenic clones. B) Denoised copy ratio scatterplots for clones 1 and 2. C) IC50 curves for clones 1 and 2 under treatment with the MCL1 inhibitor, S63845. D) IC50 curves for clones 1 and 2 under treatment with the PI3K inhibitor, AZD8186.

**Single-cell RNA-sequencing of MM cell lines with Amp1q before and after treatment with MCL1i, PI3Ki, and combination demonstrates mechanism of action** To gain insight into the mechanism of action of MCL1i, PI3Ki, and their combination in MM with Amp1q, we performed single-cell RNA sequencing on two MM cell lines with Amp1q (KMS12BM, KMS11) treated with MCL1i alone, PI3Ki alone, and their combination, compared to DMSO (**Figure 6A**). We found that both monotherapies and the combination were significantly cytostatic (i.e., increased the proportion of cells in G1), while MCL1i and the combination appeared to cause a G2M arrest with a significant increase in the proportion of cells in the S phase and a concomitant decrease in the proportion of cells in the G2M phase (**Figure 6B, C**). At the gene expression level, key genes that are important for the G2M transition, including *PLK1*, *CDK1*, and *CDC20*, were consistently downregulated in the MCL1i and combination arms (**Figure 6D**). While MCL1 has been shown to regulate the cell cycle in recent years^73–78^, it was previously thought that MCL1 inhibition with S63845 did not disrupt that functionality^79^. In the PI3Ki arm, we observed consistent downregulation of the heat-shock protein-encoding genes, *HSPA1B*, *HSPH1*, and *DNAJB1*, which are known to be regulated by PI3Ki and are important in mediating protein folding and stabilization under stress, cellular functions that we found are increasingly essential in MM with Amp1q (**Figure 6E**)^80–82^. Furthermore, we observed consistent downregulation of genes important for microtubule function, such as *TUBG*, *KIF5B*, and *ARPC5L*, which potentially explains the observed cytostatic effect and suggests that the effect of PI3K inhibition on cell cycle regulation is complementary to that of MCL1 inhibition. Notably, we observed consistent upregulation of *BCL2*, which however did not translate into increased BCL2 dependency, as shown by dynamic BH3 profiling in a MM cell line with Amp1q (**Figure 6F**).

**Figure 6.**
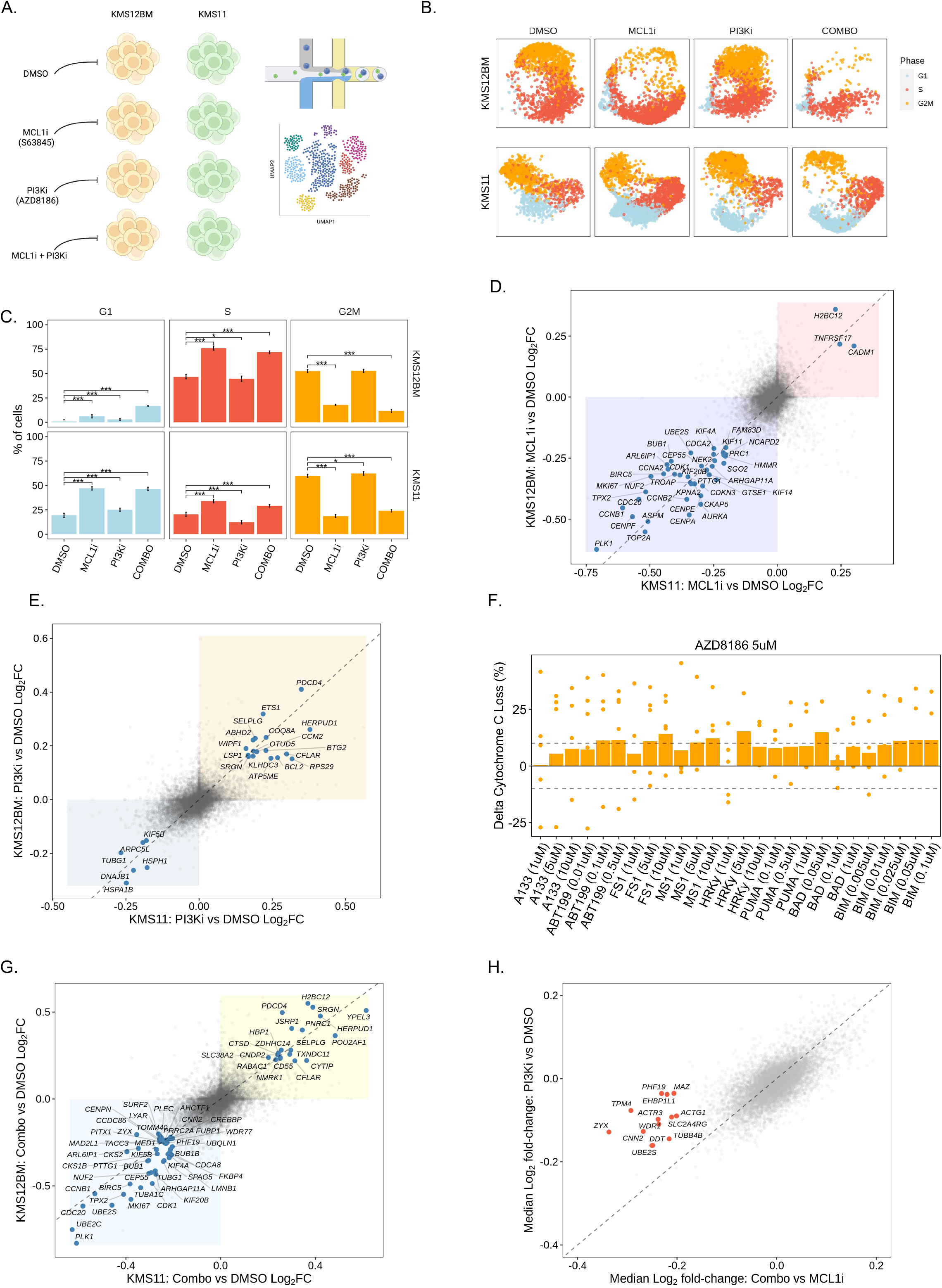
Single-cell RNA-sequencing of MM cell lines with Amp1q before and after treatment with MCL1i, PI3Ki, and combination demonstrates mechanism of action. A) Schematic of the experimental approach. B) UMAP embedding of KMS12BM and KMS11 cells, colored by cell cycle phase (G1: blue; S: red; G2M: orange) split by condition (DMSO, MCL1i, PI3Ki, COMBO). C) Barplots of cell cycle phase frequency per cell line and condition. The height of each bar corresponds to the average frequency across 10K iterations of subsampling 70% of cells; the error bar corresponds to the 95% confidence interval. P-values were computed using bootstrapping with 10K iterations. D) Scatterplot of log_2_ fold-change in expression with MCL1 inhibitor S63845 compared to DMSO in KMS11 (x-axis) and KMS12BM (y-axis). Genes with fold-changes < -0.2 or > 0.2 in both cell lines were labeled. E) Scatterplot of log_2_ fold-change in expression with PI3K inhibitor AZD8186 compared to DMSO in KMS11 (x-axis) and KMS12BM (y-axis). Genes with fold-changes < -0.15 or > 0.15 in both cell lines were labeled. F) Barplot of the difference (delta) in cytochrome *c* loss (%) between treatment with the PI3K inhibitor AZD8186 (5uM) and DMSO per peptide. Dashed lines were drawn at 10% and -10%. G) Scatterplot of log_2_ fold-change in expression with combination treatment with the MCL1 inhibitor S63845 and the PI3K inhibitor AZD8186 compared to DMSO in KMS11 (x-axis) and KMS12BM (y-axis). Genes with fold-changes < -0.2 or > 0.2 in both cell lines were labeled. H) Scatterplot of the median log_2_ fold-change in expression with combination treatment with the MCL1 inhibitor S63845 and the PI3K inhibitor AZD8186 compared to S63845 alone in both KMS11 and KMS12BM (x-axis) and median log_2_ fold-change in expression with the PI3K inhibitor AZD8186 compared to DMSO in both KMS11 and KMS12BM (y-axis). Genes with a fold-change of < -0.2 in the Combo vs MCL1i arm and > -0.2 fold-change in the PI3Ki vs DMSO arm (i.e., genes whose expression changes in Combo compared to MCL1i but past what is expected based on the activity of PI3Ki) were colored in red and labeled.

Lastly, in the combination arm, we observed a combination of the changes seen in the monotherapy arms (**Figure 6G**). Interestingly, genes associated with genome organization and cell motility, such as *ACTG1*, *MAZ*, and *CNN2*, were downregulated compared to MCL1i alone past what would be expected due to PI3Ki alone (**Figure 6H**).

## DISCUSSION

While targeted therapy can improve outcomes, the genetic landscape of MM is highly heterogeneous and does not offer itself for precision medicine, with few exceptions, most notably that of patients carrying a t(11;14) translocation^4, 19–21, 82^. Gain or amplification of chr1q provides a golden opportunity for the development of targeted therapy strategies, on account of its high frequency, its role in disease progression, and its negative impact on survival^26–45^. However, developing targeted therapy for patients with MM and Amp1q is challenging, due to: (i) the lack of isogenic cell line models, (ii) the scarcity of MM cell lines without Amp1q, and (iii) the sheer number of genes that are perturbed by Amp1q in *cis* or in *trans*^28^. Prior efforts have focused on genes located on chr1q, such as *MCL1*, *BCL9*, *CKS1B*, *ILF2*, and *PBX1*^46–56^.

Here, we have performed a large-scale systematic characterization of genetic dependencies and therapeutic vulnerabilities associated with Amp1q and identified a broad range of opportunities for targeted therapy. To reduce technical noise and control for irrelevant inter-tumor variability in our dependency analyses, we performed a genome-wide CRISPR screen and a targeted shRNA screen of genes located in 1q21-1q23, and filtered for genes that hit in multiple datasets and were differentially expressed between MM cell lines with and without Amp1q. This approach led to minimal positional bias across the genome, while enriching for hits associated with Amp1q, as demonstrated by the fact that 1q21-1q23 was the region with the highest density of genetic dependencies across the genome in MM cell lines with Amp1q. We observed increased dependency on cell cycle, metabolism, protein folding and degradation, and the PI3K-mTOR pathway. For genes located on chr1q, we confirmed the increased dependency on *MCL1*, *BCL9*, and *ILF2*, and identified novel targets, including *UAP1* and *UCHL5*, an N-glycosylation enzyme and a deubiquitylating enzyme respectively that protect cancer cells from endoplasmic reticulum (ER) stress, *CLK2*, a kinase that regulates RNA splicing, *POLR3C*, a gene encoding a subunit of RNA polymerase III which is crucial for protein biosynthesis, *PIP5K1A*, a kinase responsible for generating PIP2, the substrate of PI3K kinases, and *ZNF687*, a zinc-finger protein that activated the PI3K pathway^83–88^. Interestingly, we found that *MCL1*, *CLK2*, and *ZNF687* dependencies may be associated with Amp1q in a lineage-agnostic manner, as we showed increased dependency on those in breast and lung cancer cell lines with Amp1q.

As genetic dependencies do not always translate into actionable therapeutic vulnerabilities, we complemented our dependency screens with two drug screens. Both drug screens identified the PI3K-mTOR pathway as the top therapeutic vulnerability in MM cells lines with Amp1q, which is consistent with the results of our dependency analyses. We confirmed the increased vulnerability of MM with Amp1q to PI3K and MCL1 inhibitors in an expanded panel of cell lines and demonstrated synergistic activity between the MCL1i S63845 and the PI3Ki AZD8186 in MM cell lines with Amp1q. To confirm that the sensitivity to MCL1i and PI3Ki is indeed associated with Amp1q and not a secondary alteration that may co-segregate with Amp1q in our cell line cohorts, we performed single-cell RNA sequencing in 6 bone marrow and circulating tumor cell samples from patients with MGUS/SMM and subclonal Amp1q. The presence of a subclonal Amp1q allowed us to compare gene expression levels between cells with and without Amp1q within the same tumor, which inherently accounts for inter-tumor and inter-individual variability. Indeed, we showed that subclones with Amp1q show consistently higher expression levels of MCL1 and the PI3K-mTOR pathway. Moreover, though, we were able to identify dependencies on genes not located on chr1q, which once again revealed increased dependency on the PI3K pathway. This finding suggests that the dependency on the PI3K-mTOR pathway may be regulated both in *cis* and in *trans*. To explore what region on chr1q is responsible for conferring sensitivity to MCL1i and PI3Ki, we isolated isogenic clones from the KMS12BM MM cell line, with either 4 or 3 copies for the segment between cytobands 1q31.1 and 1q44 and compared their sensitivity to S63845 and AZD8186 *in vitro*. We showed a significant increase in sensitivity to both inhibitors in Clone 1, which has 4 copies for the entire arm. This suggests that the breadth of the amplification matters for response, and that the copy number of the 1q21-1q25 segment alone, as routinely tested clinically via FISH, may not be appropriate for patient selection.

Finally, we performed single-cell RNA sequencing on two MM cell lines with Amp1q, KMS12BM and KMS11, to characterize the mechanism of action of MCL1i and PI3Ki, as well as their combination, in MM with Amp1q. We observed a cytostatic effect with both inhibitors, although through different mechanisms: MCL1i appeared to cause a G2M block/S phase arrest, while PI3Ki appeared to interfere with microtubule function at the gene expression level. Previous studies have identified MCL1 as the key regulator of apoptosis in mitotic arrest, and a regulator of the cell cycle in general^73–78^. However, MCL1 inhibition by S63845 was previously shown to spare this functionality in HEK293T cells, where no effect was observed on either the cell cycle or apoptosis^79^. On the other hand, PI3K has been shown to play a role in microtubule stabilization, a function that can be disrupted via PI3K inhibition, but also the G1/S transition, which potentially explain the observed cytostatic effect^90–94^. Furthermore, PI3K inhibition was shown to downregulate the expression of genes encoding members of the heat-shock protein family, which play a key role in the unfolded protein response and MM cell survival^80–82^. Given the increased dependency on the unfolded protein response pathway we observed in MM cell lines with Amp1q, this effect may partly explain the therapeutic efficacy observed with PI3Ki. Lastly, in accordance with our *in vitro* assays showing synergy between MCL1i and PI3Ki in MM cell lines with Amp1q, we observed a combination of the individual gene expression effects induced by monotherapy in cell lines treated with the combination, with genes related to genome organization and cell motility showing potentially enhanced effects compared to monotherapy. Synergistic activity between MCL1 and PI3K inhibition was also recently demonstrated in breast cancer, where they observed rapid induction of apoptosis with the combination^72^.

As both MCL1 and PI3K inhibitors are toxic at therapeutic doses, the observed synergistic activity and the increased sensitivity of MM tumors with Amp1q to these inhibitors may enable their use at lower doses without sacrificing efficacy. Furthermore, as both types of inhibitors have been clinically tested, the translation of these findings will be accelerated. Overall, these findings bring us closer to establishing targeted therapy for patients with MM and Amp1q, who have not benefited as much from recent advances in MM therapeutics and are in need of better options.

## ACKNOWLEDGEMENT

The authors would like to thank the patients who participated in the PCROWD study. Additionally, the authors would like to thank Anna V. Justis, PhD, for medical editing support, and Sarah Nersersian, MSc, for illustration support. This research was supported in part by the National Institutes of Health (R35CA2263817-01A1). R.S.P. is supported by the MMRF Research Fellowship Award, the International Waldenstrom’s Macroglobulinemia Foundation’s Robert A. Kyle Award, the Claudia Adams-Barr Award for Innovative Basic Cancer Research, the Hellenic Society of Hematology, and the International Myeloma Society. G.G. is partially supported by the Paul C. Zamecnik Chair in Oncology at Massachusetts General Hospital Cancer Center. M.S.D. is supported by the National Institutes of Health (R01CA266298-01A1). S.J.F.C is supported by the Simeon J. Fortin Charitable Foundation Postdoctoral Fellowship award.

## INCLUSION AND DIVERSITY

One or more of the authors of this paper self-identifies as an underrepresented ethnic and/or gender minority in science. One or more of the authors of this paper self-identifies as a member of the LGBTQIA+ community. We support inclusive, diverse, and equitable conduct of research.

## SUPPLEMENTAL TABLES

**Supplemental Table 1.** Top 150 upregulated and downregulated genes associated with Amp1q in patients with MM, BRCA, and LUSC, used to query the Connectivity Map.

